# CIGB-300 reverses chemo-resistance in MDR1-transfected lung squamous cancer cells

**DOI:** 10.1101/2024.10.24.620006

**Authors:** Meifeng Wang, Dongfang Tang, Xiaofang Luo, Wan Liu, Jiale Xie, Ying Yi, Yaqin Lan, Wen Li, Silvio E. Perea, Wubliker Dessie, Yasser Perera, Zuodong Qin

**Affiliations:** Hunan Engineering Technology Research Center for Comprehensive Development and Utilization of Biomass Resources, Hunan University of Science and Engineering, Yongzhou 425199, China; China-Cuba Biotechnology Joint Innovation Center (CCBJIC), Yongzhou Development and Construction Investment Co., Ltd. (YDCI), Yangjiaqiao Street, Lengshuitan District, Yongzhou City, Hunan Province, 425000, People’s Republic of China; Molecular Oncology Group, Department of Pharmaceuticals, Biomedical Research Division, Center for Genetic Engineering & Biotechnology (CIGB), Havana 10600, Cuba

**Author notes:** **Corresponding authors:** Zuodong Qin; Xiaofang Luo, Yasser Perera.

**Keywords:** CIGB-300, Drug resistance, Resistance reversal, Overexpression, ABCB1

## Abstract

**Background:** Inhibition of ABC transporter protein activity is considered to be the most effective method to reverse multidrug resistance (MDR). In this study, we evaluated the MDR reversal potential of CIGB-300, a potent CK2 kinase inhibitor.

**Methods:** ABCB1 overexpressing lung adenocarcinoma NCI-H226 cells were constructed using lentivirus, and the expression of ABCB1 gene and protein was detected by real-time fluorescence quantitative PCR and Western blotting. MTT assay was used to assess the cytotoxicity and MDR reversal effect of CIGB-300.The effect of CIGB-300 on ABCB1 expression was determined by Western Blotting. Cell surface expression and subcellular localization of ABCB1 were examined by Flow Cytometry and Immunofluorescence Staining. Rh123 efflux and accumulation were measured by Fluorescent Enzyme Labeler and Flow Cytometry.

**Results:** CIGB-300 significantly increased the sensitivity of drug-resistant cells overexpressing the ABCB1 drug efflux pump (NCI-H226-ABCB1), while it had no effect on their parental cell lines. At the same time, its mechanism of action may be related to the inhibition of ABCB1 expression, which was dose-dependent, Moreover, in addition, we demonstrated that CIGB-300 reduced the expression of NFKB and CDC37 proteins.

**Conclusions:** Our study elucidated that CIGB-300 reverses ABCB1-mediated MDR by inhibiting ABCB1 protein expression or intracellular signaling and provides a potential therapeutic strategy to improve tumor chemosensitivity.

## 1. Introduction

Cancer is a serious disease that is one of the greatest threats to human health, with lung cancer topping the list of all malignant tumors^[1]^. Chemotherapy is the most common treatment for cancer. However, in the treatment of cancer, multidrug resistance (MDR) is a common problem that poses a challenge to the medical community, due to its ability to hinder the efficacy of chemotherapy. Tumor multidrug resistance occurs by a variety of mechanisms and involves a number of factors; one of the main reasons for this is the increased expression of the ATP binding cassette (ABC) family of drug efflux transporters^[2-5]^. ABC transporter proteins are structurally highly conserved, and they can decrease or abolish the effect of chemotherapy by coupling ATP binding and hydrolysis to reduce the intracellular accumulation of drugs to sub-therapeutic levels^[6-8]^. To date, at least 15 ABC transporter proteins have been associated with cancer chemo-resistance, and one of them in particular, P-glycoprotein (P-gp, ABCB1), is overexpressed in various cancers and excretes a wide range of chemotherapeutic agents, making it an attractive therapeutic target^[9-11]^.

The most direct way to overcome multidrug resistance caused by ABC transporter proteins is to block or modulate their activity^[12]^. To date, four generations of ABC transporter protein modulators have been developed, including natural compounds, tyrosine kinase inhibitors and small molecules^[13-22]^. Unfortunately, their clinical trials have had only limited success, mainly due to lack of specificity and low safety profile ^[23-25]^. These limitations have prompted efforts to find safe and effective transporter protein inhibitors.

In recent years, casein kinase CK2 has emerged as a viable tumor target and different kinase inhibitors have been developed, including small molecule ATP competitors, synthetic peptides, and antisense oligonucleotides ^[26-28]^. Our previous study showed that CIGB-300, a novel anticancer peptide, impaired CK2-mediated phosphorylation by directly binding to conserved phosphoreceptor sites on the substrate and significantly modulated the expression levels of drug resistance-associated proteins and genes^[29-31]^. However, the relationship between CIGB-300 and drug-resistant lung cancer cells is unclear, while its role in MDR1-mediated drug-resistant lung cancer cells has not been reported. In this study, we establish an ABCB1 overexpressing lung cancer cell line, H226/ABCB1, and evaluate the MDR reversal potential of CIGB-300 to discover new indications for this compound in preventing chemotherapy resistance in cancer patients.

## 2. Materials and methods

### 2.1 Chemicals and reagents

CIGB-300 was synthesized as described before^[32]^; 1-(4, 5-dimethylthiazol-2-yl)- 3, 5-diphenylformazan (MTT), verapamil, docetaxel and other chemicals were purchased from Sigma-Aldrich (St. Louis, MO, USA). RPMI 1640 medium was obtained from Thermo Fisher Scientific Inc. (Waltham, MA, USA). Antibodies against ABCB1/P-gp (K112141P), HDAC1 (K001617P), p-HDAC1 (K010474P), NFKB (K101551P), p-NFKB (K009559P), CDC37 (K110309P), p-CDC37 (K010475M), HMGB1 (K108830P) and Glyceraldehyde-3-phosphate dehydrogenase (GAPDH) were obtained from Solarbio Life Science Co. (Beijing, China). ReverTra Ace® qPCR RT Kit and SYBR Green qPCR Master Mix were purchased from TOYOBO Bio (Shanghai, China).

### 2.2 Cell cultures

H226-PURO and NCI-H226-ABCB1 cells were established by transfecting wild-type NCI-H226 [H226] (ATCC CRL-5826) with either the empty pHBLV-CMV-MCS-3flag-EF1-puro (H226-PURO) DNA or vector containing the full length ABCB1 (NCI-H226-ABCB1), and were cultured in a medium containing 0.5 μg/mL of puromycin. All cells were cultured in flasks with RPMI 1640 supplemented with 10% (v/v) fetal bovine serum and 1% (v/v) penicillin/streptomycin at 37℃ in a humidified incubator containing 5% CO_2_. The cells were harvested using 0.25% trypsin once they reached 80% confluence.

### 2.3 Lentivirus package and stable cell lines construction

The overexpression lentivirus vector of MDR1 and the knockdown lentivirus vector of MDR1 were constructed and packaged by Hanbio Co., Ltd., named as HBLV-h-ABCB1-P and HBLV-PURO, respectively. For constructing ABCB1 stable overexpression cells, NSCLC cell line NCI-H226 was infected with HBLV-h-ABCB1-PURO virus or control virus in the presence of 7.0 µg/ml Polybrene. 48 h later, cells were selected with 1.0 µg/ml puromycin for 3 weeks. Further qRT-PCR was used to examine the overexpression efficiency of HBLV-h-ABCB1-PURO in cells^[33]^.

### 2.4 Cell viability assay

The viability of H226-puro and H226-ABCB1 cells to anticancer drugs was measured using the MTT assay. Cells (6000 cells/well) were seeded in 96-well plates (100 μL/well) and cultured for 24 h. For the cytotoxicity experiment of CIGB-300, 100 μL of varying concentrations of CIGB-300 were added into each well to the final concentrations of: 250 μM, 166 μM, 111 μM, 75 μM, 49 μM, and 33 μM. They were incubated for 48 h at 37 ℃. For the reversal experiment, CIGB-300 (50 μL/well) was added 1 h prior. 50 µl of different concentrations of chemotherapeutic drug (docetaxel) were then added to the indicated wells and incubated for 48 hours at 37 ℃. Subsequently, 20 μL of the MTT solution (5 mg/mL) were added to each well and incubated for an additional 4 h. The solution was discarded and 100 μL of DMSO were added to dissolve the formazan crystals. Finally, light absorbance was determined at 562 nm using a multi-plate reader (Filter Max F3, Molecular Devices, San Jose, CA, USA)^[34, 35]^.

### 2.5 Quantitative real-time PCR (qRT-PCR)

Total RNA from cells was extracted with Trizol. Complementary DNA (cDNA) was then reversely transcribed with a Rever Tra Ace® qPCR RT Kit according to the standard procedure. ABCB1 levels were examined with a Light Cycler 96 Real-Time Fluorescence PCR (Roche, Switzerland). GAPDH served as an internal reference. The 2-^△△ct^ method was performed to calculate the relative expression of ABCB1.

### 2.6 Western blot analysis

To determine whether CIGB-300 affects the protein expression of ABCB1 and CK2 substrates, after cells were harvested after pre-treatment and incubation with CIGB-300 for 6 h or 24 h, they were harvested and lysed with RIPA lysis buffer for 20 min on ice. The protein lysate was then measured with a BCA protein assay kit, and an equal amount of denatured protein lysate was separated by SDS-PAGE and transferred onto 0.45µm polyvinylidene fluoride. The membrane was blocked with PBS with 5% nonfat milk for 2 h and incubated with primary antibody at 4℃ overnight, followed by incubation with a second antibody for 1 h. The immunoblots were visualized by an enhanced chemiluminescence ECL kit. Cropping and densitometric analysis of Western blot images were performed using Image J^[36, 37]^. Target protein levels were normalized to the indicated reference proteins.

### 2.7 Flow Cytometry

Cells in the logarithmic growth phase were seeded in 6-well plates with 2.5 × 10^5^ cells/well and treated for 48 h. The cells were then collected and washed 3 times with PBS at 4°C, and subsequently 2 μL of ABCB1 antibody were added to them, while a second tube was left untreated. Afterwards, samples were incubated at room temperature for 60 min in a dark chamber. The measurements were performed on a Beckman flow cytometer, and the data were analyzed using FlowJo software^[37]^.

### 2.8 Immunofluorescence Staining

Approximately 1×10^5^ cells were seeded on glass coverslips and incubated for 24 h at 37℃. The medium was then removed, the cells were washed with PBS, and incubated with fresh medium containing IC_20_ concentrations of CIGB-300 for another 48 h. PBS treatment served as a control. Subsequently, cells were fixed with 4% paraformaldehyde for 20 min, permeated with 3% hydrogen peroxide, blocked with 5% bovine serum albumin for 2 hat room temperature, and incubated with polyclonal antibodies against ABCB1 at 4℃ overnight. After washing, the cells were incubated with an Alexa Fluor® 488 Rabbit Anti-Goat IgG for 1 h and stained with DAPI for 5 min at room temperature. Images were collected using a laser scanning confocal microscope (Zeiss, LSM800, Zeiss Corporation, Germany)^[36, 38]^.

### 2.9 Rh123 accumulation assay

The effect of CIGB-300 on the intracellular accumulation of Rh123 was measured by flow cytometry as previously described. Briefly, cells were incubated with different concentrations of CIGB-300 at 37 °C for 24 h. 1μg/ml Rh123 was added and cells were cultured for another 30min. Finally, the cells were digested and collected, washed 3 times with chilled phosphate-buffered saline (PBS) and analyzed by flow cytometry (CytoFLex, Beckman Coulter Inc., Brea, CA, USA)^[39, 40]^.

### 2.10 Rh123 efflux assay

Rh123 efflux assay was performed as previously described^[39, 40]^. NCI-H226-ABCB1 cells were treated with different concentrations of CIGB-300 at 37 °C for 6 h. Rh123 at 1 μg/ml was then added and the cells cultivated for another 30min. After removal of CIGB-300 and 3 washing with PBS, fresh medium was added to the cells to measure the extrusion of Rh123. Finally, the cells were collected and lysed at indicated time points (0, 60, 90, 120 and 180 min), and Rh123 fluorescence intensity both in the culture medium and cellular lysates were analyzed by Fluorescent Enzyme Labeler (Biotek 800 TS, USA).

### 2.11 PI staining assay

Cells were incubated with 40μM, 74μM and 100μM CIGB-300 during 6h; then replaced with R10 incubated another 120min, and suspended in 100µl binding buffer (1X). Next, 5 µl of Propidium Iodide (PI) were added and incubated in dark for an additional 15 min. Finally, cells were added with 400µl binding buffer(1X) and subsequently analyzed by Flow Cytometry (Becton-Dickinson BD, USA) with the acquisition of 15000 events/sample. The data analysis was performed using the FlowJo software v10.

### 2.12 Statistical analysis

All data were obtained from at least three independent experiments (n ≥ 3), Statistical analysis was performed using the software SPSS 25.0 and. Data results were expressed as mean ± S.D (standard deviation), a t-test was used for intergroup comparisons, and one-way ANOVA was used for multiple mean comparisons; P<0.05 was considered statistically significant for differences of results.

## 3. Results

### 3.1 NCI-H226-ABCB1 model construction and chemo-sensitivity testing

Firstly, we constructed the cellular model NCI-H226-ABCB1 cells with stable over-expression of the corresponding multi-drug resistant gene via lentivirus infection of NCI-H226 lung squamous carcinoma cells (LUSC). The expression of ABCB1 in such target cells were subsequently evidenced at gene (qPCR) and protein (Flow Cytometry, Western blot) level. qPCR results indicated that ABCB1 expression was significantly increased in NCI-H226-ABCB1 but not vector-transduced NCI-H226-PURO cells (Fig.1A). Additionally, Western blot analysis confirmed the over-expression of ABCB1 in NCI-H226-ABCB1 cells (Fig.1B). Finally, flow cytometry confirmed that ABCB1 pump levels at the surface of live NCI-H226-ABCB1 cells was least 8-fold higher than on its NCI-H226-PURO counterpart (MFI ratios=8x) (Fig.1C).

**Fig. 1:**
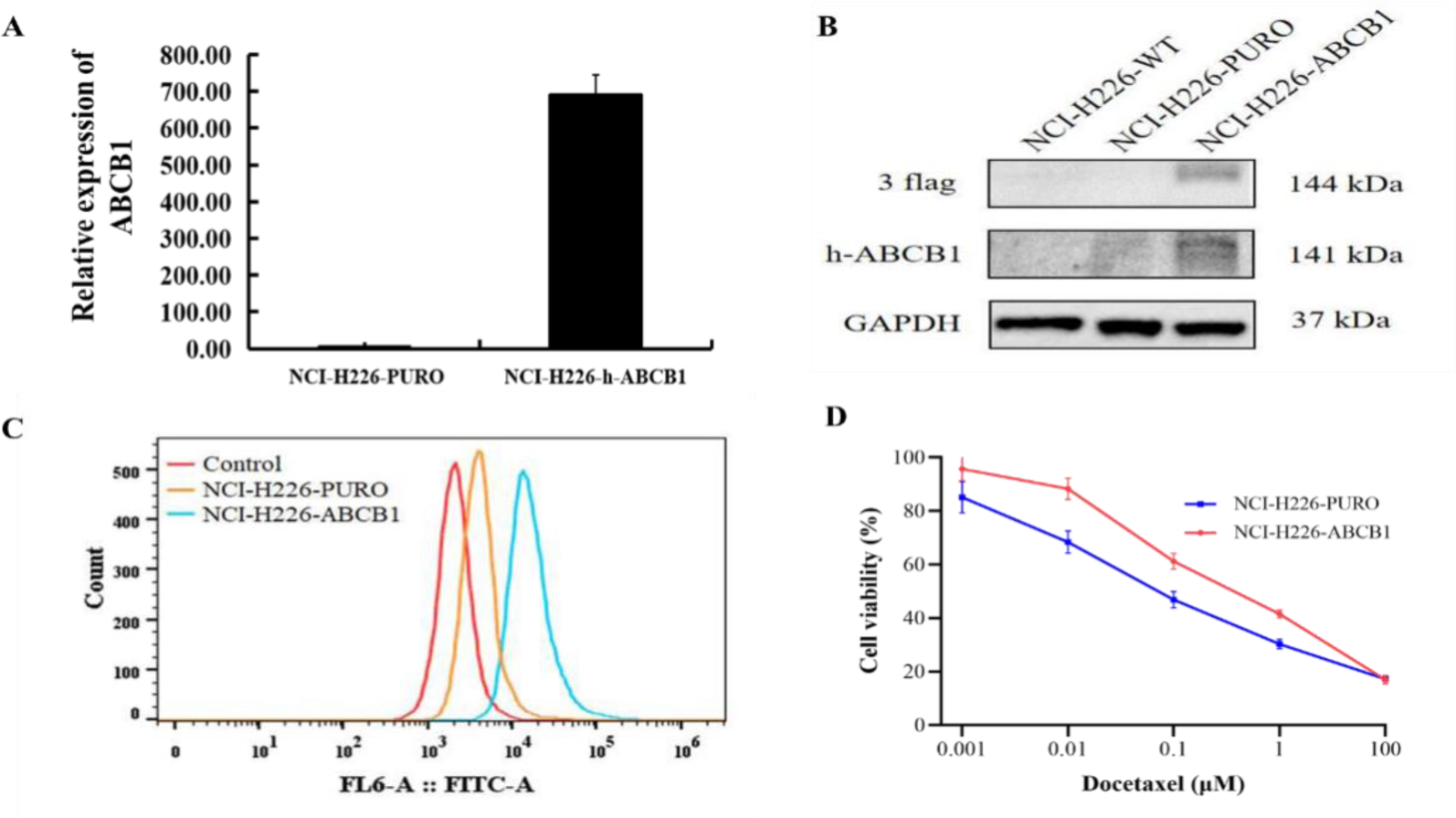
Detection of ABCB1 mRNA expression, corresponding total/surface protein expression and chemo-resistance to docetaxel chemotherapy in NCI-H226-PURO and NCI-H226-ABCB1 cellular models. Quantification of mRNA levels and protein expression by RT-PCR (A), Western blotting (B), or flow cytometry on live cells (C). Evaluation of cytotoxicity by MTT (D). Data points represent the means ± SD of at least three independent experiments performed in triplicate.

Since ABCB1 pump plays a critical role in docetaxel resistance, we subsequently measured the chemo-sensitivity of NCI-H226-ABCB1 cells by the MTT method. Fig.1D shows the typical dose-response curves, yet on NCI-H226-ABCB1 cells, cell survival after incubation with increasing concentrations of docetaxel was higher than in their vector infected counterparts (ie., NCI-H226-PURO cells). Accordingly, the IC_50_ or potency of docetaxel in this drug-resistant cells was 0.658 μM; whereas, in NCI-H226-PURO cells the IC_50_ value was 0.095 μM, thus indicating around 7-fold lower potency of docetaxel for cell killing in NCI-H226-ABCB1 drug resistant cells.

### 3.2 CIGB-300 restore docetaxel chemo-sensitivity in NCI-H226-ABCB1 cells

The structure of the synthetic peptide chimera CIGB-300 is shown in Fig. 2A. The polycationic residues of the Cell Penetrating Peptide (CPP) Tat are well exposed to the solvent and mediate the internalization of the inhibitory p15 cyclic domain within the low micro-molar range^[41]^. To select optimal peptide concentrations for drug sensitization experiments, firstly we evaluate the concentration-dependent cytotoxic curves for CIGB-300 on NCI-H226-ABCB1 and NCI-H226-PURO cells (Fig. 2B). The results indicated that NCI-H226-ABCB1 cells display the same sensitivity to CIGB-300 than their NCI-H226-PURO counterparts; thus, indicating a no drug cross-resistance effect in this cellular model.

**Fig. 2:**
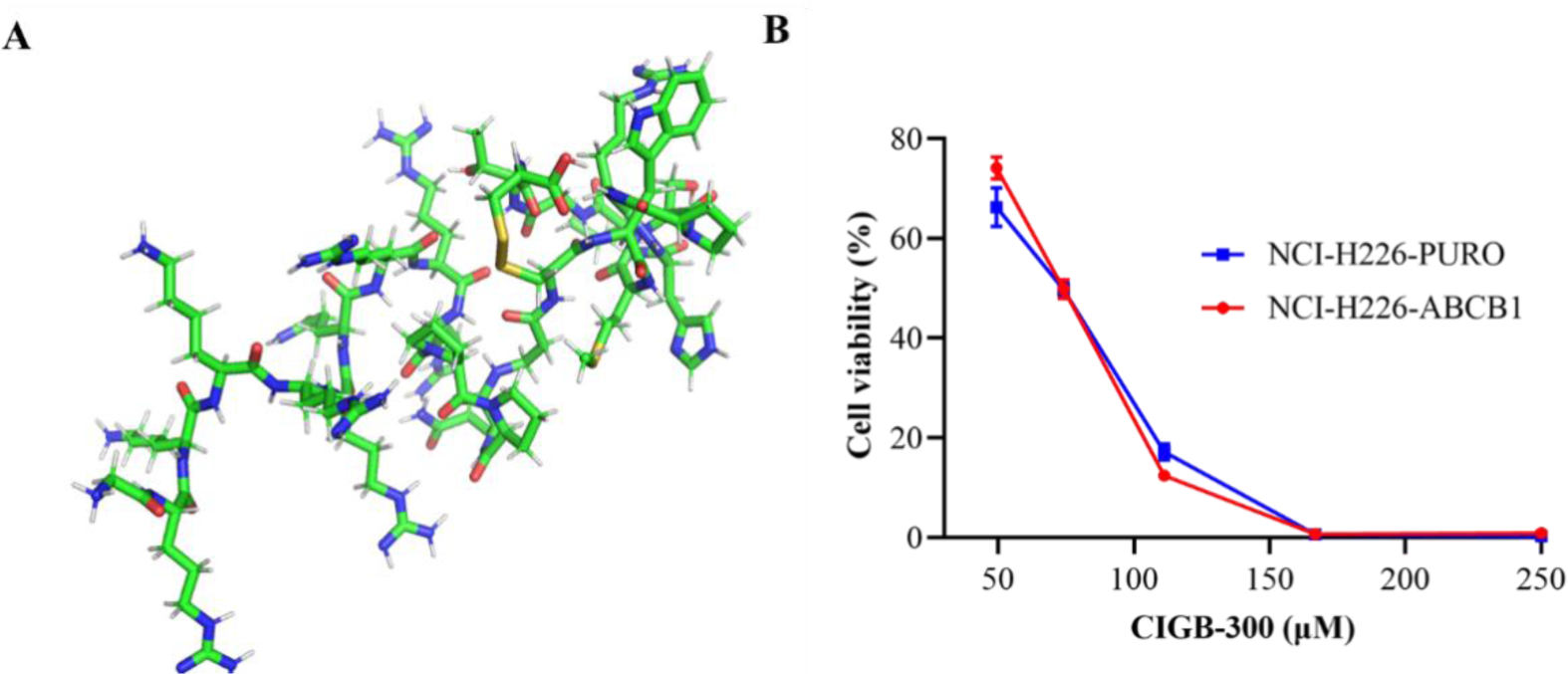
Modeled structure and cytotoxicity effects of CIGB-300 by MTT assay. (A), the structure of CIGB-300 was modeled by Pymol; (B), MTT cytotoxicity assay was used to measure cell survival in NCI-H226-PURO and ABCB1-overexpressing NCI-H226-ABCB1 cells. All cells were exposed to CIGB-300 for 48 h. Data points represent the means ± SD of at least three independent experiments performed in triplicate.

According to resulting cytotoxicity curves, more than 80% of cell survival rate is registered below 40 μM of CIGB-300 concentrations. Therefore, we selected sub-optimal cell killing concentrations of 20, 30 and 40 μM of CIGB-300 for subsequent chemo-resistance reversal experiments upon incubation with docetaxel.

To evaluate the chemo-sensitivity to docetaxel of NCI-H226-ABCB1 cells when co-incubate with sub-optimal doses of CIGB-300 we performed MTT assays. Table 1 compares the calculated IC_50_ values from the dose-response curves of docetaxel co-incubated with CIGB-300 or vehicle in both NCI-H226-ABCB1 and NCI-H226-PURO cells. As expected, NCI-H226-ABCB1 cells exhibited higher resistance to the ABCB1 substrate docetaxel than parental cells^[42]^. However, the addition of CIGB-300 to docetaxel partially reverse the acquired resistant phenotype at peptide concentrations of 20 μM, it increased the potency back around 3x from 0.622 to 0.216 uM (2.87-fold); whereas, had little effects on the survival of NCI-H226-PURO cells, but still pre-sensitizes the cancer cells for killing (IC_50_ of docetaxel going down).

**Table 1:**
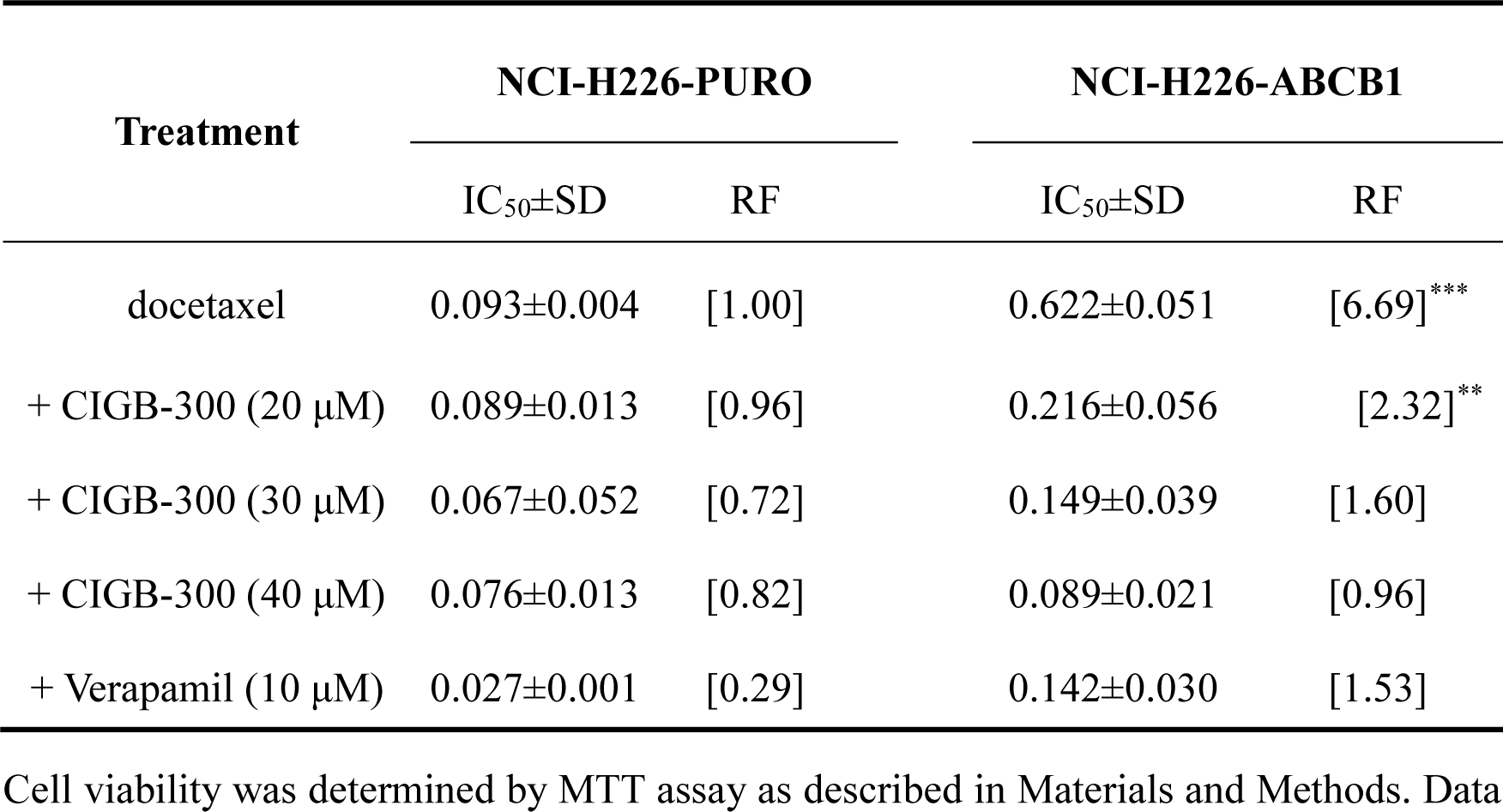
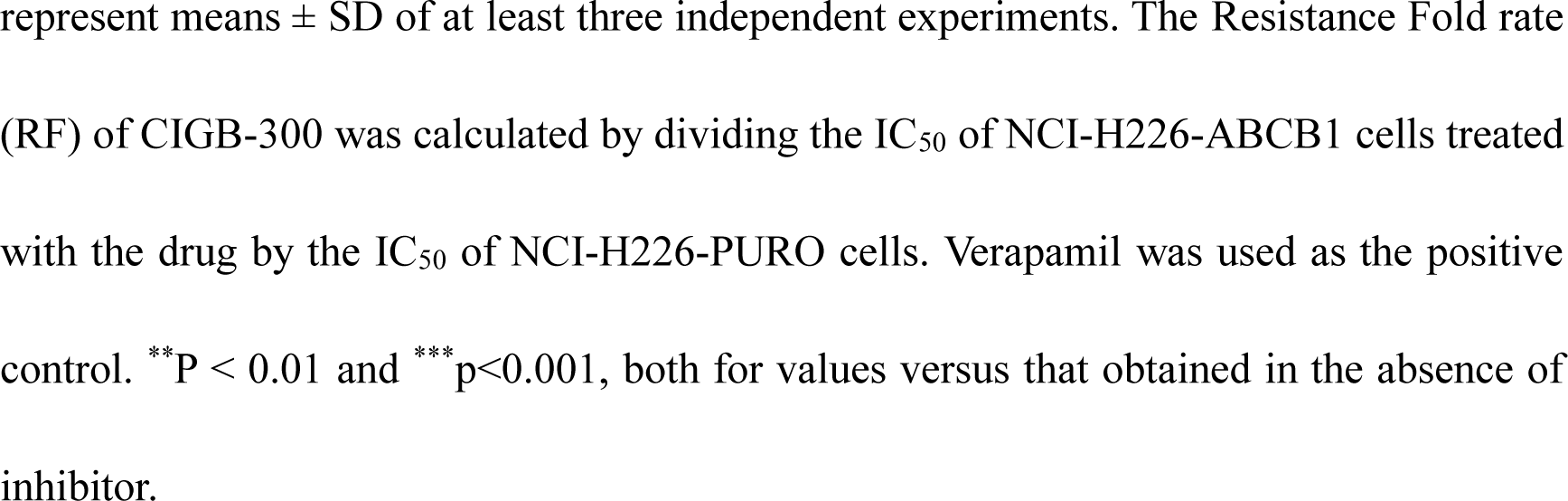
Chemo-sensitivity analysis of CIGB-300 combined with docetaxel affecting H226-ABCB1 cells.

| Treatment | NCI-H226-PURO |  | NCI-H226-ABCB1 |  |
| --- | --- | --- | --- | --- |
| | IC<sub>50</sub>±SD | RF | IC<sub>50</sub>±SD | RF |
| docetaxel | 0.093±0.004 | [1.00] | 0.622±0.051 | [6.69]<sup>***</sup> |
| + CIGB-300 (20 μM) | 0.089±0.013 | [0.96] | 0.216±0.056 | [2.32]<sup>**</sup> |
| + CIGB-300 (30 μM) | 0.067±0.052 | [0.72] | 0.149±0.039 | [1.60] |
| + CIGB-300 (40 μM) | 0.076±0.013 | [0.82] | 0.089±0.021 | [0.96] |
| + Verapamil (10 μM) | 0.027±0.001 | [0.29] | 0.142±0.030 | [1.53] |

Cell viability was determined by MTT assay as described in Materials and Methods. Data
represent means ± SD of at least three independent experiments. The Resistance Fold rate (RF) of CIGB-300 was calculated by dividing the IC<sub>50</sub> of NCI-H226-ABCB1 cells treated with the drug by the IC<sub>50</sub> of NCI-H226-PURO cells. Verapamil was used as the positive control. \*\*P < 0.01 and \*\*\*p<0.001, both for values versus that obtained in the absence of inhibitor.

### 3.3 Effect of CIGB-300 on ABCB1 expression in NCI-H226-ABCB1 cells

The above results indicated that CIGB-300 could effectively sensitize MDR cells with high/ectopic ABCB1 expression to chemotherapeutic drugs like docetaxel. This finding could be explained by down-regulation of the corresponding transporter protein^[43, 44]^; thus, subsequently we analyzed the effect of CIGB-300 on ABCB1 protein expression by immunofluorescence, western blot and flow cytometry (Fig. 3).

**Fig. 3:**
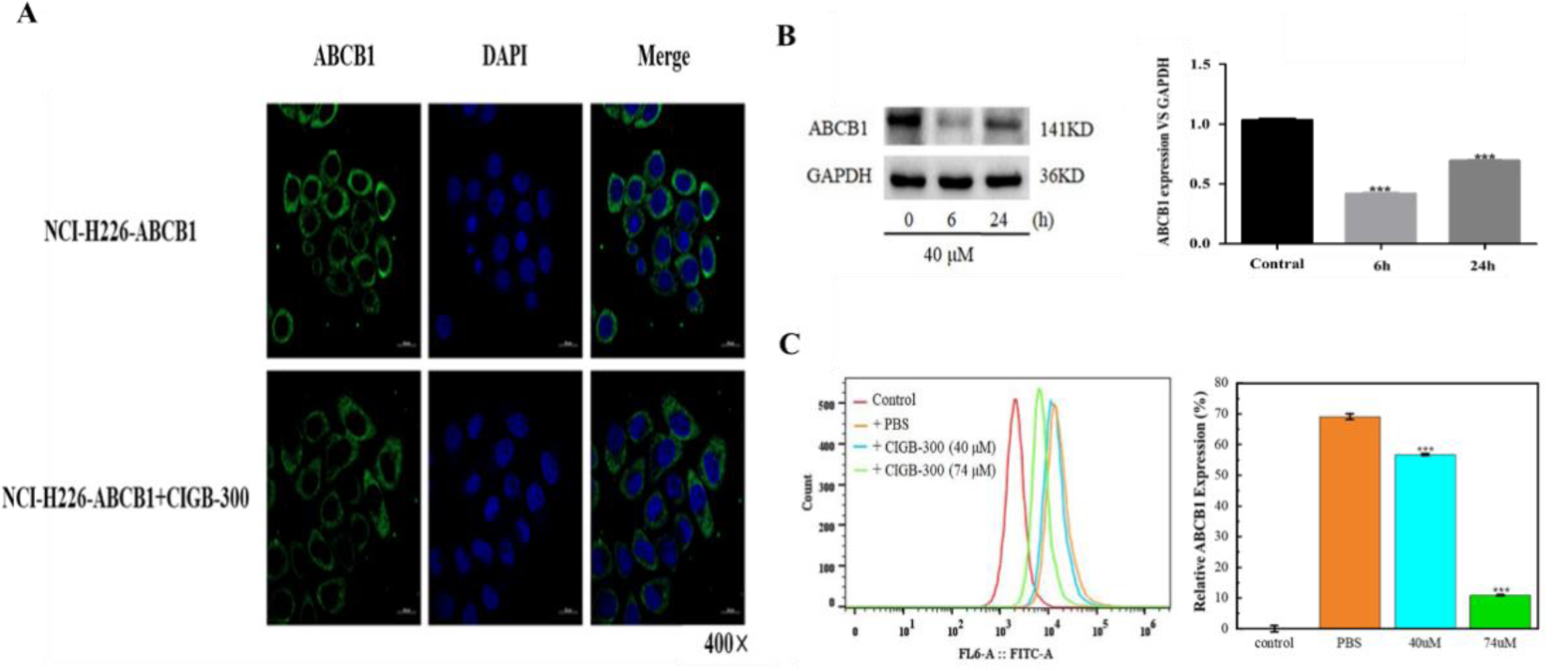
Effect of CIGB-300 on the ABCB1 protein expression in NCI-H226-ABCB1 cells. (A), immunofluorescence staining after 48 h incubation with 40 μM of peptide; (B), Western blotting for ABCB1 detection in Total Cell Extracts. GAPDH was used for protein load normalization; (C), flow cytometry after 48 h incubation with 40 or 74 μM of peptide. Data points represent the means ± SD of at least three independent experiments performed in triplicate, ***p<0.001, both for values versus that obtained in the absence of inhibitor.

After incubating NCI-H226-ABCB1 cells with 40 μM CIGB-300 for 48 h, the fluorescence intensity of ABCB1 became weaker and its expression was reduced compared to the control. Meanwhile, the ABCB1 levels in Total Cell Extracts (TCE) also evidenced a clear reduction of ABCB1 protein, with a peak decreasing at 6 h and some sort of recovery at 24 h (ie., transient effect). In addition, when explored two different concentrations of CIGB-300 at a fixed time of 48 h, we observed a dose-dependent effect on ABCB1 protein levels in the surface of intact NCI-H226-ABCB1 cells.

### 3.4 Effect of CIGB-300 on ABCB1 function in NCI-H226-ABCB1 cells

Multidrug resistance reversal agents can also function by impair pump activity of ABCB1 transports. Thus, we investigated the effect of CIGB-300 on ABCB1 functional activity at different measurement times. Unexpectedly, when NCI-H226-ABCB1 cells were treated with two different concentrations of CIGB-300, the extracellular level of Rh123 was increased, while the intracellular accumulation of Rh123 was decreased (Fig. 4). Of note, CIGB-300 penetrates/permeabilizes cellular membranes as part of its mechanism of action; thus, we further evaluated whether this result could be explained by a partial permeabilization of NCI-H226-ABCB1 cells by CIGB-300. As noted in Fig. 4B, even 40 μM of CIGB-300 concentration promotes PI entry into the cells, such permeabilization of the cancer cells was more evident at 74 μM, thus explaining the observed increase in the extracellular levels of Rh123 (eg., 90min time-point, Fig.4A). Therefore, the observed Rh123 release upon CIGB-300 incubation seemed unrelated to ABCB1 pump over-activation.

**Fig. 4:**
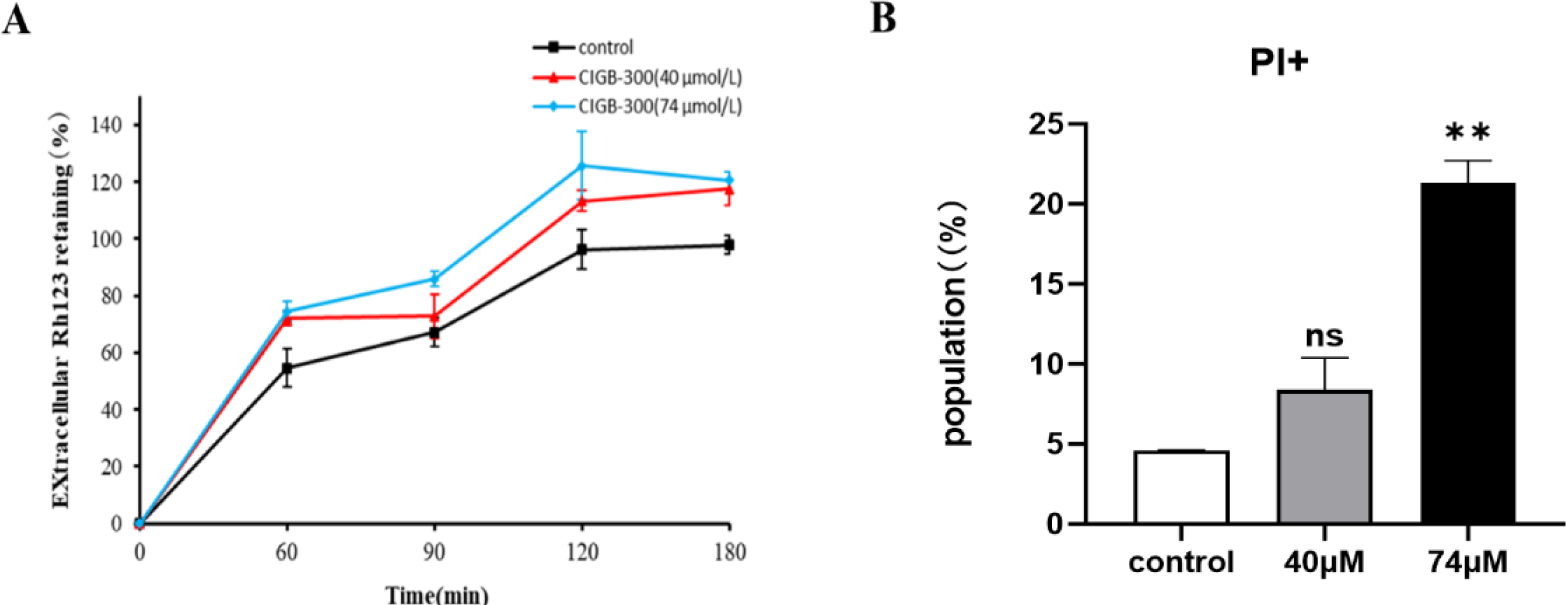
Effect of CIGB-300 on the ABCB1 function measured as efflux of Rh123 to the cell supernatant and remained intracellular content. (A), efflux of Rh123 after 6h of incubation with two concentrations of CIGB-300 at a fixed time point of 180 min; B) PI staining experiments on the NCI-H226-ABCB1 cell line treated with CIGB-300 during 6h. Here, PI+ cell populations indicate the PI penetration into the cells. Data points represent the means ± SD, ***p<0.001,

### 3.5 Exploration of CK2 substrates related with chemo-resistance of cancer cells

Besides ABCB1 expression/activity, several CK2 substrates have been related to chemo-resistance in cancer cells^[45]^. Therefore, we measured by western blot the effect of CIGB-300 over the expression of CK2 associated proteins in NCI-H226-ABCB1 cells. Fig. 5 shows that after incubation of NCI-H226-ABCB1 cells with CIGB-300 at 40 μM the expressions of HDAC1, P-HDAC1 (55 KDa), CDC37 and P-CDC37 (44 kDa) were increased, while the expressions of NFKB (32 KDa) and P-NFKB (35 kDa) were decreased compared to vehicle (PBS).

**Fig. 5.**
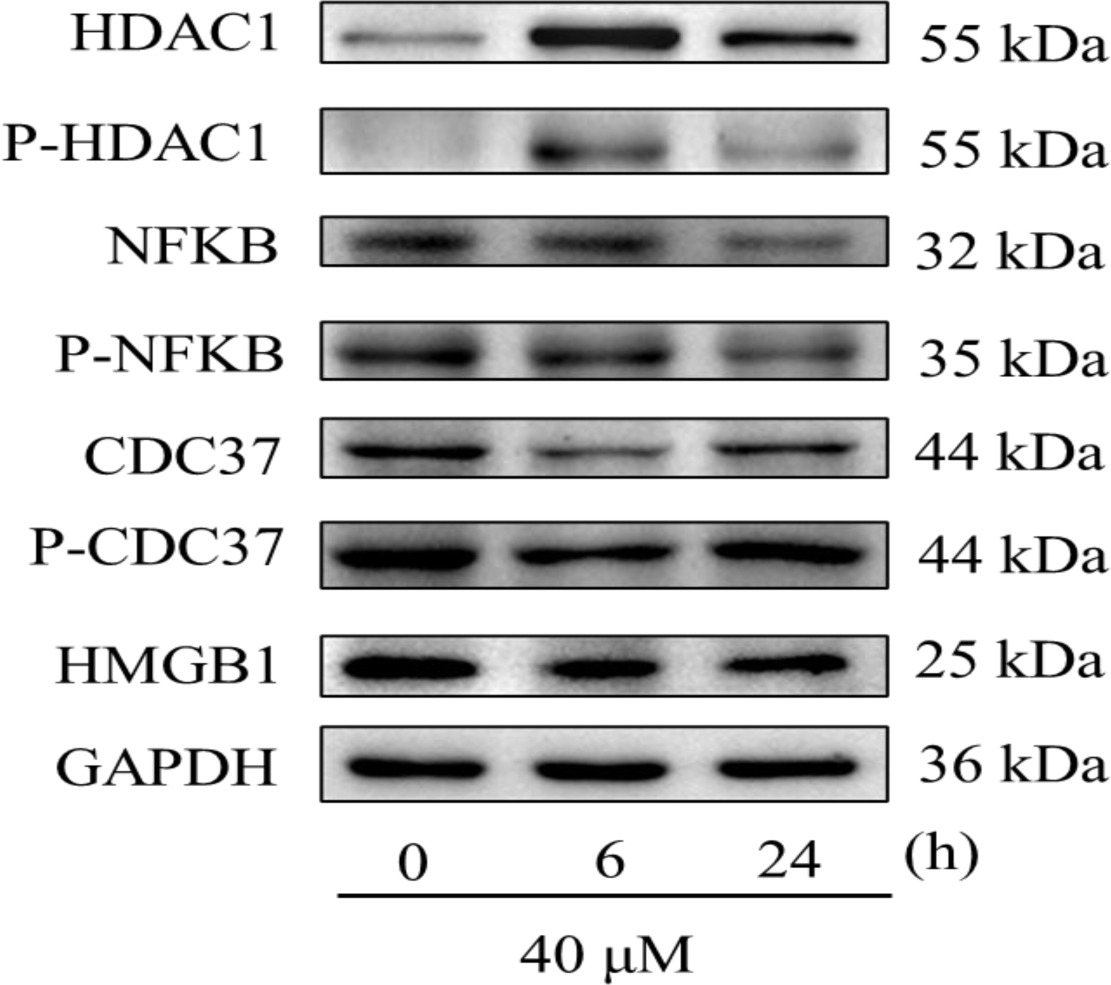
Western blot analysis of selected CK2 substrates in TCE from NCI-H226-ABCB1 cells incubated with CIGB-300. Data points represent the means ± SD of at least three independent experiments performed in triplicate.

## 4. Discussion

The development of safe and effective multidrug resistance reversal agents remains clinically unsatisfactory. Acquired multidrug resistance, including over-expression of some ABC transporter proteins, is one of the main reasons for chemotherapy failure^[46]^. To date, three generations of ABCB1 inhibitors have been tested in clinical trials; unfortunately, however, clinical trials have not yielded satisfactory results overall. Meanwhile, research on the development of fourth-generation inhibitors, including natural active compounds and peptide drugs, is still underway^[47-49]^.

CIGB-300 is a clinical grade anti-protein kinase CK2 peptide that binds to the phosphorylated receptor site on CK2 substrates and to CK2α catalytic subunit^[41]^. We have previously reported CIGB-300 inhibitory effects on tumor cell lines derived from breast, lung, and leukemia cancers, and it can be used after surgery to limit the metastatic spread of tumors^[50-52]^. However, studies on CIGB-300 antineoplastic effects on chemo-resistant lung cancer cells, as well as its purported capacity to reverse multidrug resistance are limited^[51]^. One previous study showed that CIGB-300 effectively inhibited the proliferation of ciplatin-resistant A549 lung cancer cells^[49]^.

In this study, we found that a sub-optimal concentration of CIGB-300 enhanced the therapeutic effect of docetaxel when co-incubated in NCI-H226 ectopically expressing ABCB1 protein, thus providing first evidences of partial reversion of drug resistance in a cellular model of LUSC.

The *in vitro* modulatory potency of CIGB-300 was investigated for by MTT assay. The cytotoxic activity of CIGB-300 was assayed *in vitro* with NCI-H226 cancer cell lines stably expressing ABCB1 (Fig. 2B), and an *in vitro* reversal assay was performed using suboptimal concentrations of CIGB-300 for cell killing (<40 μM) (Table 1).

To study the mechanism by which CIGB-300 may reverse the induced ABCB1-mediated multidrug resistance, the effect of the peptide over total and surface ABCB1 protein expression were examined. Our results showed that in the presence of CIGB-300, both ABCB1 total and cell surface expression are reduced (Fig. 3).

In addition to MDR1 total and cell surface level reductions upon incubation with CIGB-300, the reversal of MDR1 phenotype may be achieved by affecting the functional activity of ABCB1^[34]^. However, instead of reduction of Rh123 extrusion to cell supernatant, we detected some increase in extracellular levels of Rh123 from NCI-H226/ABCB1 treated cells. These findings contradict the observed sharp down-regulation on ABCB1 levels upon incubation with CIGB-300, as well as the subsequent sensitization of NCI-H226-ABCB1 to docetaxel. By performing PI staining of NCI-H226-ABCB1 incubated cells, we demonstrated that even at 40 μM the CIGB-300 affect the cell membrane permeability and this may cause an artefactual exiting of Rh123 (ie. not mediated by MDR1 pumping activities). Altogether, we demonstrated that CIGB-300 may decrease overall levels of ABCB1 resistant-related drug efflux transporter and this may explain at least partially the reversion of resistant phenotype to docetaxel.

Another potential mechanism for chemo-resistance reversal could be associated with the impairment of phosphorylation of several CK2 substrates (eg., NFKB, CDC37) previously associated with drug resistant in cancer cells^[45]^. To evaluate this possibility, we performed western blot experiments in TCE from cells incubated with CIGB-300. Our data showed no significant changes in high mobility group box-1 protein (HMGB1) expression, decreased expression of nuclear factor-kappa B (NFKB) and cell division cycle 37 (CDC37) and their phosphorylated proteins, yet increased expression of histone deacetylase 1 (HDAC1) and its phosphorylated proteins in the NCI-H226-ABCB1 cell line (Fig. 5). Consistent with our previous study, CIGB-300 can provide new insights into the mechanism of action of CIGB-300 by inhibiting the CK2-dependent pathway and thus reversing drug resistance in cancer cells^[51]^.

Taken together, the present study demonstrated that CIGB-300 was able to reverse ABCB1-mediated MDR *in vitro*. The mechanism of MDR reversal by CIGB-300 involves inhibition of ABCB1 expression, but probably also derived from the impact of intracellular signaling related to general chemo-resistance like NFKB and CDC37 pathways. These results suggest that CIGB-300 may have potential as a reversal agent, which in combination with conventional anticancer drugs could re-enforce the sensitivity of tumor chemotherapy.

## 5. Conclusions

CIGB-300 reverses multi-drug resistance by selectively and effectively inhibiting ABCB1 protein expression in one model cancer cells of LUSC origin. These findings provide strong support for further development of CIGB-300 or its derivatives as candidate inhibitors of MDR in cancer therapy.

## Ethics approval and consent to participate

Not applicable.

## Consent for publication

All of the authors mutually agree for submitting this manuscript to the Journal of Experimental & Clinical Cancer Research.

## Availability of data and material

Not applicable.

## Authors’ contributions

**Meifeng Wang**: Conceptualization, Methodology, Investigation, Actual experimentation, Writing – review & editing, Writing – original draft, **Dongfang Tang, Xiaofang Luo, Wan Liu, Jiale Xie, Ying Yi, Yaqin Lan, Silvio E. Perea,** and **Wubliker Dessie**: Observation recording, **Wen Li, Yasser Perera,** and **Zuodong Qin**: Supervision, Project administration.

